# Genome Analyses of A Worldwide Multisource Collection of *Klebsiella variicola* Reveal Adaptation Relevant for Human Infection

**DOI:** 10.1101/2024.10.28.617863

**Authors:** Jiayu Wang, Claudia E. Coipan, Eelco Franz, Ed Kuiper, Els Wessels, David L. Gally, Karthick Vasudevan, Balaji Veerarhagavan, Refath Farzana, Nicola Holden, Ruth Zadoks, Paul G. Higgins, Thamarai Schneiders

## Abstract

*Klebsiella variicola* (*K*.*var*), part of the *Klebsiella pneumoniae* (*K*.*pne*) species complex, is an emerging human pathogen originally associated with plants. Phylogenomic analyses of a global dataset of isolates categorised as human, animal, plant, or environmental strains suggest that *K*.*var* are broadly disseminated regardless of hosts or habitats and exhibit genomic variability that is relevant for occupying and transmitting within and across multiple sources. Ancestral state analyses confirm that *K*.*var* are originally derived from non-human sources. Genome-wide association analyses of human versus non-human isolates of *K*.*var* indicate isolates from human origins are linked to the loss of cold-shock response genes such as *lpxP* and *cspB*. Importantly, human *K*.*var* isolates also showed enriched antimicrobial resistance gene content and diversity acquired through mobile genetic elements, horizontal gene transfer and mutations within the chromosomal loci which underscore the impact of antimicrobial exposure and proximity to antibiotic resistant *K*.*pne* in nosocomial environments. Capsular profiles of *K*.*var* suggest high levels of variability in gene composition and indicate that gene functionality for *wcaJ* in capsule formation differs between *K*.*var* and *K*.*pne*. Although capsular specialisation was not observed in a host/habitat or lineage-dependent manner, human isolates harbour, on average, a higher siderophore gene content in addition to *mrkD* mutations which impact on fimbriae and biofilm formation. There is limited evidence of any specific *K*.*var* lineages exhibiting convergence of both virulence and antimicrobial resistance genotypes which suggest that these attributes are evolving independently in clinical strains. Our work highlights that gene attrition and acquisition provide a basis for adaptation of *K*.*var* as human pathogens. The broad distribution and evolution of *K*.*var* highlight the need for improved diagnostic precision and surveillance under a One Health framework.

## 1 Introduction

*Klebsiella variicola* (*K*.*var*), a member of the *Klebsiella pneumoniae* (*K*.*pne*) species complex (KpSC), is an emerging human pathogen [1, 2, 3]. Previously associated with plants [4, 5], animals [6, 7], and environmental sources [8, 9], more recent studies using whole-genome sequencing (WGS) indicate an increasing prevalence in human infections [3, 2, 1, 10].

The recent and growing recognition of *K*.*var* as a human pathogen is partially due to the overlapping biochemical profiles of *K*.*var* and *K*.*pne*, which have resulted in the misidentification of *K*.*var* using standard laboratory detection methodologies [11]. Whilst this may contribute to the lower detection rates, epidemiological surveillance using whole-genome sequencing rectifies this misclassification [12] and reveals the true prevalence of *K*.*var* as a human pathogen, whereby current estimates suggest that 2.5% to 10% of isolates that are identified as *K*.*pne* in clinical laboratories are actually *K*.*var* [11, 13, 14].

Like the more prevalent *K*.*pne, K*.*var* can cause human infections including neonatal sepsis, pneumonia, bloodstream, and urinary tract infections [15, 1, 3]. Intriguingly, *K*.*var* has been linked to higher rates of mortality associated with bloodstream infections in adults and neonates [2, 10], in addition to increased bladder burden and urovirulence [1] relative to *K*.*pne*. However, the exact microbial factors that drive these higher mortality rates in bloodstream infections and bacterial burden in urinary tract infections remain uncharacterised.

Importantly, *K*.*var* has been linked to disease in other hosts such as ironwood trees [5], dairy cattle [7], and commensalism as endophytes [16] and symbiotes [17] in plants. Recent microbiome analyses of bovine urinary samples [18] and the human gut [19, 20] identified *K*.*var* strains which further highlight its role as part of the endemic microbiota.

This wide-ranging lifestyle, from commensalism to virulence, attests to opportunism and suggests genome plasticity in adaptation to different hosts and/or habitats. Previous studies [1, 21, 8] using smaller datasets underscore a lack of restriction between different *K*.*var* genomic lineages in a host-, habitat-, or geographic locale-dependent manner, which posits that ubiquitous gene acquisition and transmission events across different and within the same sources remain a distinct possibility. In support of this, current data show that horizontal gene transfer of the capsule locus can be acquired from *K*.*pne* through horizontal gene transfer in nosocomial environments [3, 1, 22].

Therefore, the broad dissemination and aptitude for gene exchange with *K*.*pne* position *K*.*var* as an important emerging human pathogen. However, the adaptive traits that facilitate the survival and dissemination of *K*.*var* in different hosts or habitats remain unknown. Thus, in this study, using the largest genomic dataset to date, we sought to address these traits and their implications for human infection.

## 2 Results

### 2.1 *Klebsiella variicola* DATABASE

We acquired 723 *Klebsiella variicola* genome sequences and identified 715 as *Klebsiella variicola* ssp. *variicola* (*K*.*var*), with an Average Nucleotide Identity (ANI) of 99.03%. The remaining 8 were identified as *Klebsiella variicola* ssp. *tropicalensis*, with an ANI of 97.01%. We focused on *Klebsiella variicola* ssp. *variicola* and collected strain-specific metadata, which included the collection year, geographic origin, and source (Supplementary Table 1). Our dataset reveals the broad distribution of *K*.*var* over 26 years (1994-2020), covering six continents and diverse sources including human, animal, plant, and environmental sources. This allowed us to group those strains as linked to human (n=441) or non-human (n=248) sources or as strains with missing host/habitat information (n=26). We identified a total of 94,553 SNPs in the core genome alignment and used it for the reconstruction of the maximum-likelihood (ML) core genome phylogeny of *K*.*var*, which reveals a star-like phylogenetic structure, originating from a single common ancestor, expanding into different hosts, habitats, and geographic locations without obvious clustering (Figure 1A). The number of unique genes reached convergence (Figure 1B), showing that our *K*.*var* database is representative of the overall *K*.*var* population. The average genome size of *K*.*var* is 5.71 Mb, where the total pan-genome comprising 42,231 genes includes 4,373 core genes (at 95% percentage identity) and 37,858 accessory genes (Figure 1C). We attempted to define the *K*.*var* population structure using existing MLST schemes, namely Kleborate [23], and the *K*.*var*-specific MLST scheme described in [21], as shown in Supplementary Figure 1.

**Figure 1.**
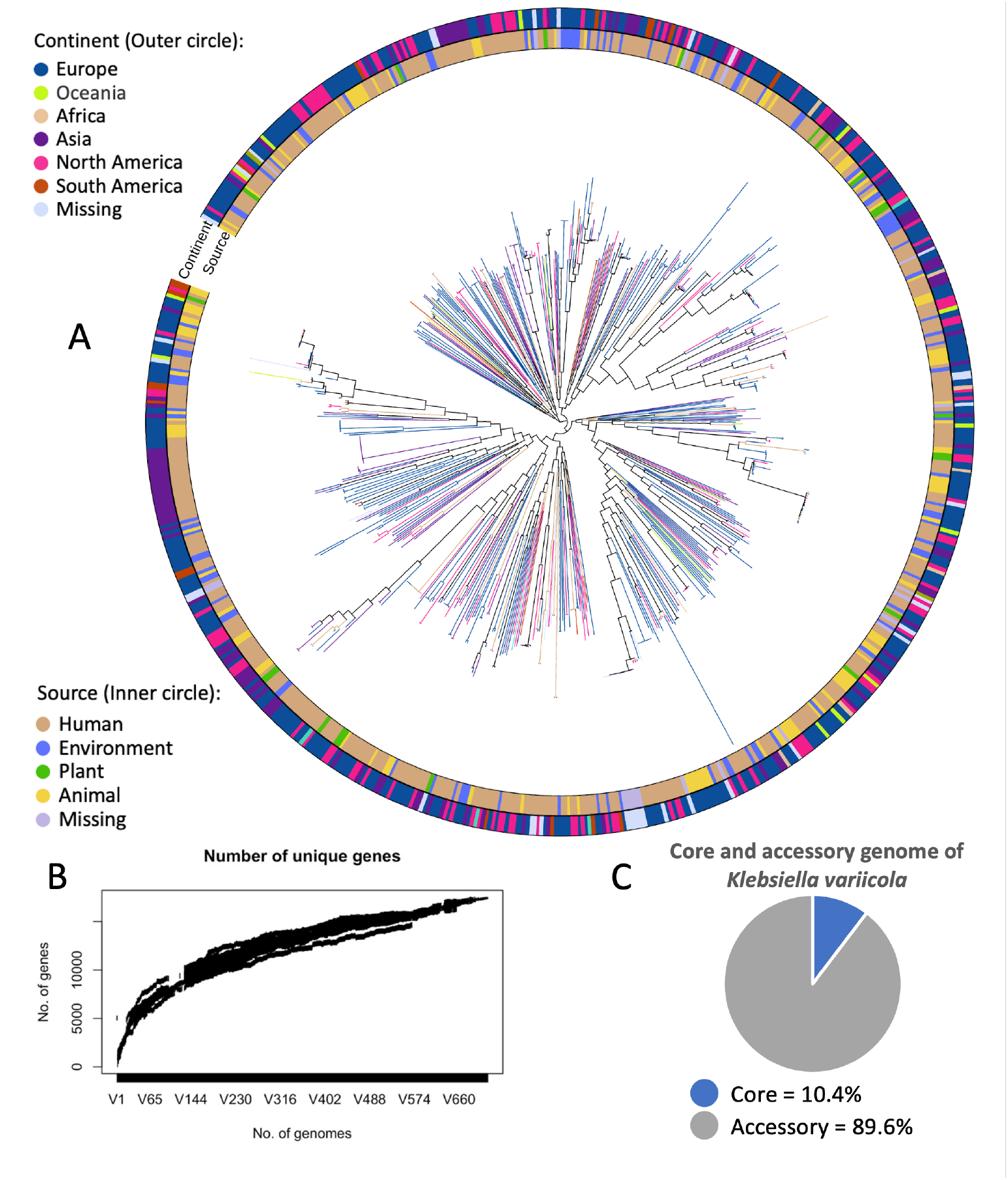
Population Structure and pan-genome of *Klebsiella variicola*. A) **Core Genome Phylogeny**: The core genome phylogeny of 715 *Klebsiella variicola* strains. Metadata annotations include host/habitat type (inner circle) and geographic origin (outer circle). The phylogenetic tree was constructed using ParSNP [25], incorporating genome data and metadata from NCBI. B) **The Number of Unique Genes**: A graphical representation plotting the number of uniquely identified genes within 715 *Klebsiella variicola* strains. C) **Pan-genome Composition**: The proportion of core and accessory genes in 715 *Klebsiella variicola* strain’s pan-genome.

### 2.2 MLST

The Kleborate MLST scheme, which was originally designed for *K*.*pne*, includes the housekeeping genes *infB, gapA, pgi, mdh, phoE, tonB*, and *rpoB*. The concordance of the Minimum Spanning Tree (MST) (Supplementary Figure 1A) with the core genome phylogeny, as indicated by Baker’s Gamma [24], was low (0.42), specifically due to allelic variations in the *K*.*var infB* gene. Among the 343 STs identified in this dataset by Kleborate MLST, 179 STs contain *infB* allelic type 24, and 103 contain *infB* allelic type 18 (Supplementary Figure 1). Taken together, these allelic types predominantly divide the *K*.*var* population into two clusters, ST347 and ST2994 (founder STs) (Supplementary Figure 1A), thus erroneously constraining the *K*.*var* MST. The *K*.*var* specific MLST scheme, MLSTKv [21], encompasses the *leuS, pgi, pgk, phoE, pyrG, rpoB*, and *fusA* genes. Using this MLST scheme against our dataset (Supplementary Figure 1B), we noted a significant shift in the distribution of founder sequence types (STs) described previously [21]. In contrast to previous reports [21, 3], the founder STs like ST10, ST38, ST23, and ST130 were replaced by novel and unassigned allelic profiles such as 1 (*leuS*), 4 (*pgi*), 3 (*pgk*), 1 (*phoE*), 11 (*pyrG*), 1 (*rpoB*), 2 (*fusA*), and 1 (*leuS*), 10 (*pgi*), 1 (*pgk*), 1 (*phoE*), 1 (*pyrG*), 1 (*rpoB*), 2 (*fusA*) (Supplementary Figure 1B). Importantly, more than 200 genomes in our study were found to have 86 novel and unassigned allelic profiles, as detailed in Supplementary Table 2. These novel allelic profiles currently lack official ST number assignments, which limits the use of the *K*.*var* schema MLSTKv [21]. Nonetheless, ST10, ST61, ST20, and ST68 remain prevalent ST-types within the larger dataset (Supplementary Figure 1B). As both approaches provided limited insights into the population structure of all *K*.*var* genomes in this dataset, we employed ParSNP [25] to construct a recombination-free, SNP-based core genome phylogeny.

### 2.3 Population structure and evolutionary dynamics

To determine the timing of *K*.*var* population expansion and transmission events, we conducted time-measured phylogenetic analyses using Beast [26] (Figure 2A). These analyses confirmed that there is limited evidence of host-or habitat-adapted lineages, indicating that *K*.*var* strains were broadly disseminated over a 1000-year time scale. Next, to trace the origins and dissemination of *K*.*var*, we utilised BayesTraits [27] for ancestral state reconstruction. The root probability analyses with the *κ* model suggest a non-human origin for ancestral *K*.*var*, with the probability of originating from a human source (0.2093) lower than that for the non-human sources (0.7907) (Figure 2B). In support of this observation, Multi-Dimensional Scaling (MDS) identifies SPARK 2777 C1, an environmental isolate collected in Pavia, Italy, between June 2017 and November 2018 [8], as the most representative *K*.*var* strain (Figure 2C and 2D). We next ascertained the rates of transition from the non-human to human source and vice versa. Our data show that transition rates from the non-human to the human source were more than double (0.3545) relative to that from human to non-human sources (0.1497), which reveals a predominance of gene flow of *K*.*var* from the non-human host/habitat into the human host.

**Figure 2.**
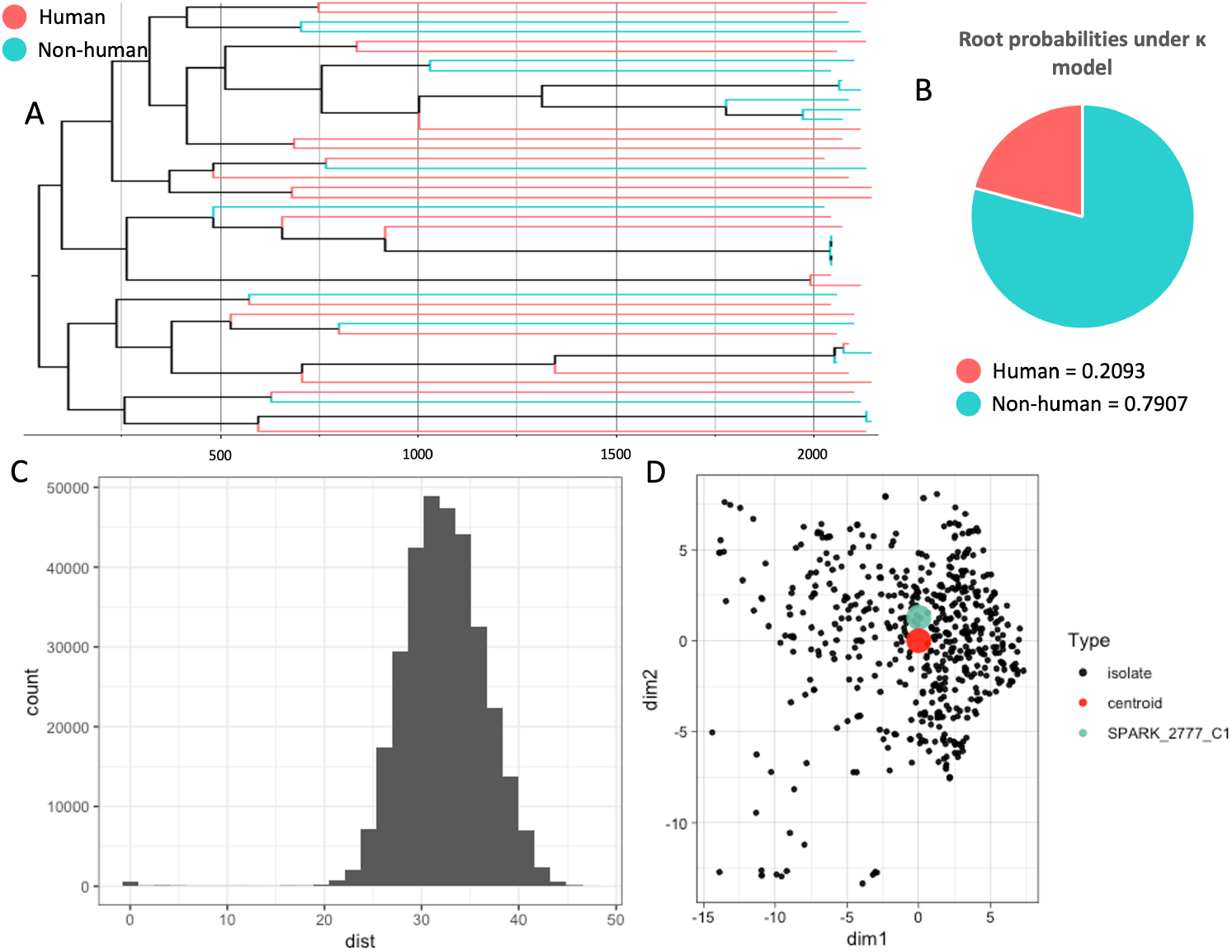
Ancestral state reconstruction and Multi-Dimensional scaling of *Klebsiella variicola*. A) **Time-Scaled Phylogeny:** A time-scaled phylogenetic tree of a subset of *Klebsiella variicola* data (n = 45) (Supplementary Table 3). B) **Ancestral State Reconstruction:** The inferred ancestral state (human or non-human) of the single common ancestor of 715 *Klebsiella variicola*. The probabilities of these ancestral states were calculated using maximum likelihood, incorporating a model that includes the *κ* parameter [27]. C) **Multi-Dimensional Scaling Distances Distribution of 715 *Klebsiella variicola* strains:** A histogram of the distribution of Multi-Dimensional Scaling (MDS) distances. The reference point at the lower left of the figure (0, 0) corresponds to the centroid of the overall data. D) **Centroid Visualization and Typing:** This panel visualizes the centroid point and 715 *Klebsiella variicola* strains in the first two dimensions. Each dot represents an isolate, with the centroid point marked in red. The pairwise Euclidean distances of each isolate from the centroid are calculated and ranked. The ‘type’ *Klebsiella variicola*, defined as the strain closest to the centroid (strain SPARK 2777 C1), is highlighted in blue.

Given its role in infection and genetic proximity to *K*.*pne* in humans, we next moved to establish the rates of genetic recombination both within *K*.*var* and between *K*.*var* and *K*.*pne*. We found that the rate of recombination within *K*.*var* populations (1.57%) is notably lower relative to that within *K*.*pne* (9.27%) alone. However, the rates of recombination between the core genomes of *K*.*var* and *K*.*pne* (8.69%) are notably higher than the rate within *K*.*var* alone, underscoring the ability for gene transfer between the two species.

### 2.4 Traits relevant to human infection

We next undertook both a selective (*nif* gene cluster) and pan-genome-based approach (GWAS) to establish whether specific genes are associated with habitat survival and adaptation.

Given that the *nif* cluster is linked to the plant- and environmental-associated lifestyle [28, 16, 4] and lost in *K. pne* potentially as a consequence of adaptation to human infections, we hypothesised that adaptation to human infection would alter the gene content of this locus. The *nif* gene cluster is usually found in the chromosome of both commensal and pathogenic diazotrophic species such as *Klebsiella* M5aI [29] and *Azotobacter vinelandii* [30], where it is responsible for synthesis, assembly, and regulation of nitrogenase levels. Nitrogenase, a key factor in plant metabolism, is the enzyme responsible for converting nitrogen (N_2_) to ammonia (NH_3_). The *nif* gene cluster consisting of core genes including *nifH, nifD*, and *nifK* encodes the nitrogenase reductase component and the *α* and *β* subunits of the nitrogenase molybdenum-iron (MoFe) protein. The *nifN, nifE*, and *nifB* genes are involved in the synthesis of the MoFe-cofactor, precursor, and the assembly and insertion of iron-sulphur clusters into the nitrogenase components respectively [28]. The *K*.*var nif* cluster, including the core genes *nifH, nifD, nifK, nifN, nifB*, and *nifE*, and accessory *nif* genes *nifA, nifF, nifJ, nifL, nifS*, and *nifW* are highly conserved, where gene phylogenies do not show any cladistic associations linked to different hosts or geographic locations (Supplementary Figure 2). The major SNPs found within the *nif* gene cluster were assessed using PolyPhen-2 and shown not to result in deleterious mutations. Consistent with our SNP analyses, the *nif* gene cluster of *K*.*var* is highly conserved regardless of host/habitat, indicating that the *nif* cluster likely plays a role that is consistent with a free-living diazotroph lifestyle.

To extend our analyses, we next applied Genome-Wide Association Analysis (GWAS) to conduct a scan of the *K*.*var* pan-genome for niche-adaptive traits. As the SNP-based core genome phylogeny did not reveal cladistic associations to host/habitat adaptation, we applied GWAS based on gene presence and absence. Using both phylogenetic-based GWAS with homologous recombination [31], and pan-genome GWAS [32] which identifies associations between gene presence/absence and other variables, e.g. habitat or host, we examined the pan-genome gene carriage differences of *K*.*var* isoltaes from human (n = 441) versus non-human (n = 248) sources.

The phylogenetic-based GWAS identified 14 candidate genes, with only 5 of these passing the correlation analysis, Fisher’s exact test, and validation through Bonferroni and False Discovery Rate (FDR) corrections. Notably, the top five genes identified in the pan-genome GWAS matched those found in the phylogenetic-based GWAS, illustrating a concordance between these two analytical methods. As shown in Table 1, the genes that are significantly associated with *K*.*var* isolates from the non-human sources are group 1575 (hypothetical protein), *cspB*, group 3155 (hypothetical protein), *chbP*, and *lpxP*. Amongst these 5 GWAS candidates, *cspB* and *lpxP* stand out in underpinning viability at lower temperatures. As shown in Table 1, *cspB* has the most significant *p* value (both Bonferroni- and FDR-corrected *p* values are *<* 0.001) and is present in 150 (60.48%) non-human associated *K*.*var* strains, while absent from 338 (76.64%) human associated *K*.*var* strains. It is a cold-shock domain protein, which plays a pivotal role in cold sensitivity at temperatures lower than 25°C by stabilising cellular processes [33]. LpxP is required for the incorporation of palmitoleolyl into lipid A under the cold-shock response, where the loss of *lpxP* results in perturbations to outer membrane integrity conferring increased sensitivity to both rifampicin and vancomycin in *E. coli* [34]. Importantly, *lpxP* is found highly conserved in those pathogens found outside the human host, further highlighting its relevance in *K*.*var* from the non-human host/habitats [35]. However, the phylogenetic analysis of these genes did not show cladistic associations with human or non-human hosts or habitats (Supplementary Figure 3).

**Table 1:** Proteins or ortholog groups from the pan-genome that are significantly associated with the non-human sources in *Klebsiella variicola* from Genome-Wide Association Analysis (GWAS). 441 human and 248 non-human *Klebsiella variicola* strains were applied in the GWAS analysis. The protein or group name and descriptions are presented in the first and second columns. The odds ratio values are reported in the third column, and Bonferroni [87] and False Discovery Rate (FDR) [86] corrected *p* values are shown in the fourth and fifth columns. The percentage of genes present in non-human strains or absent in human strains is calculated by the number of strains that carry or do not carry the gene divided by the total number of non-human (248) or human (441) strains.

| Protein/Group | Description | Odds-ratio | Bonferroni corrected $p$ value | FDR corrected $p$ value | Present in non-human strains | Absent in human strains |
| --- | --- | --- | --- | --- | --- | --- |
| group_1575 | Hypothetical protein | 5.221 | 5.00e-18 | 2.50e-18 | 150 (60.48%) | 338 (76.64%) |
| CspB | Cold-shock domain-containing protein | 5.221 | 5.00e-18 | 2.50e-18 | 150 (60.48%) | 338 (76.64%) |
| ChbP | Chitin catabolism related protein | 5.072 | 1.41e-17 | 3.52e-18 | 113 (45.56%) | 287 (65.08%) |
| group_3155 | Hypothetical protein | 5.072 | 1.41e-17 | 3.52e-18 | 113 (45.56%) | 287 (65.08%) |
| LpxP | Lipid A biosynthesis related protein | 4.983 | 3.26e-17 | 6.53e-18 | 111 (44.76%) | 288 (65.31%) |

Although GWAS analyses did not identify specific genes associated with human *K*.*var*, iNext [36] analyses showed that the overall pan-genome diversity of human associated *K*.*var* strains is higher than that in non-human associated *K*.*var* strains, where this difference is partly due to AMR gene content (shown in Figure 3A and 3B).

**Figure 3.**
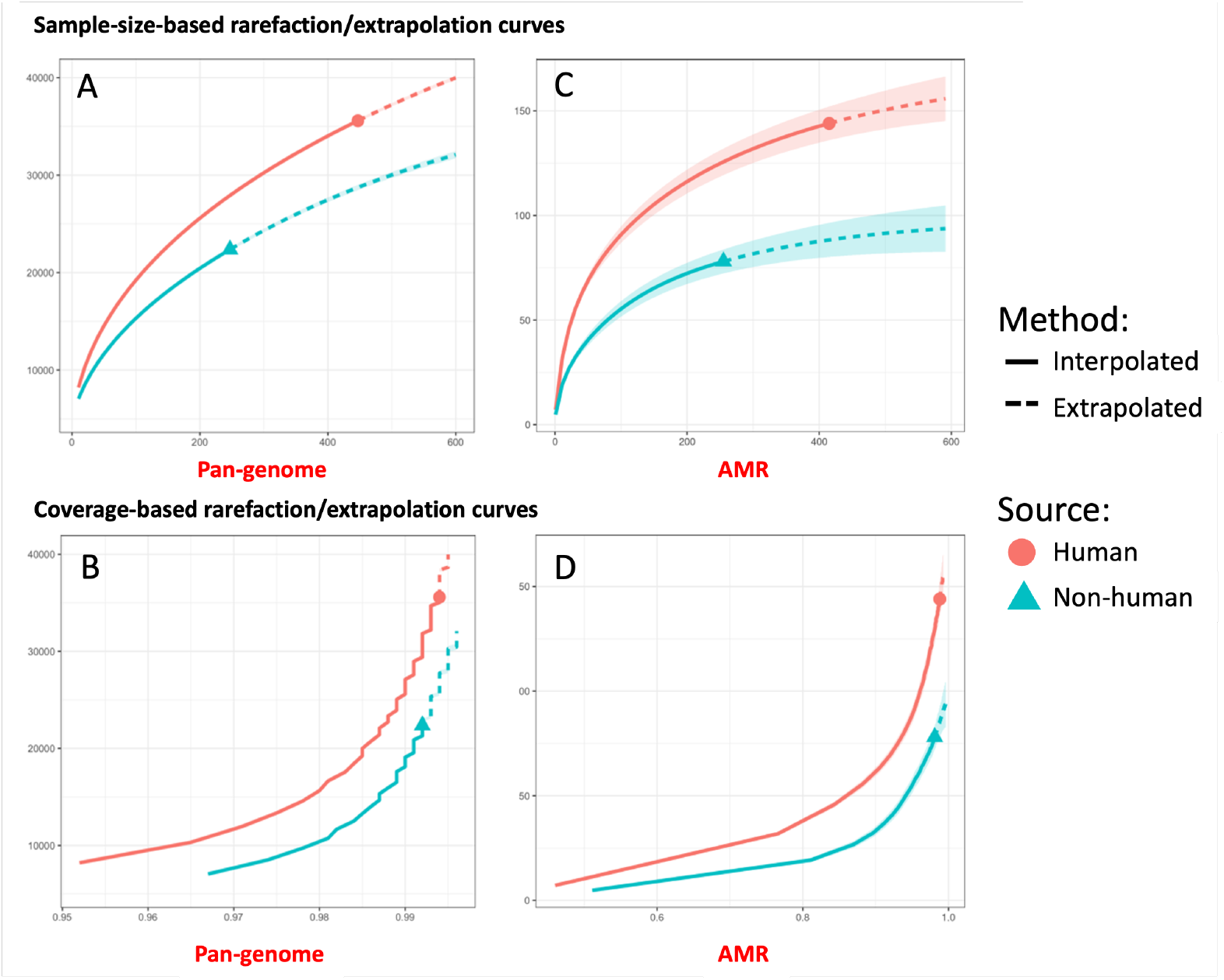
Comparative Analysis of Gene Content Diversity in Human and Non-Human *Klebsiella variicola*. iNext [36] was applied to calculate the diversity of the pan-genome and resistome of 441 human and 248 non-human *Klebsiella variicola*. A) **Pan-Genome Diversity (Sample-Size-Based Curves):** The rarefaction and extrapolation curves based on sample size for pan-genome diversity. B) **Pan-Genome Diversity (Coverage-Based Curves):** Coverage-based rarefaction and extrapolation curves. C) **AMR Diversity (Sample-Size-Based Curves):** Rarefaction and extrapolation curves for antimicrobial resistance (AMR) gene diversity, calculated based on sample size. D) **AMR Diversity (Coverage-Based Curves):** Coverage-based rarefaction and extrapolation curves for AMR gene diversity. In all panels, solid lines represent interpolated data, while dotted lines indicate extrapolated data. The human host is depicted with red lines, contrasting with blue lines that represent a combination of strains isolated from non-human hosts/habitats. Strains that do not have host/habitat annotations and the Bangladeshi isolates have been excluded from this analysis.

### 2.5 Antimicrobial Resistance

To address the variations in pan-genome diversity, we analysed the differences between human and non-human associated *K*.*var* isolates using Abricate [37]. Our analyses generally show that the *K*.*var* resistome includes genes that are both intrinsically present (core genome) e.g., *ramR, ramA, marR, acrR, oqxAB*, and *bla*_LEN_, and extrinsically acquired (accessory genome) e.g., *qnr, tet, cat, bla*_KPC_, *bla*_NDM_, and *bla*_OXA_ (Figure 4). We next compared the total AMR gene content between *K*.*var* obtained from human and non-human sources to determine if there are any significant associations with the human host. Comparisons of human (n = 441) and non-human (n = 248) *K*.*var* strains revealed that human strains (mean = 7.173) contain a significantly higher number of AMR genes (*p* value *<* 0.001) relative to non-human associated *K*.*var* strains (mean = 6.174) (Supplementary Table 4) as well as a greater diversity of AMR genes (Figure 3C and 3D).

**Figure 4.**
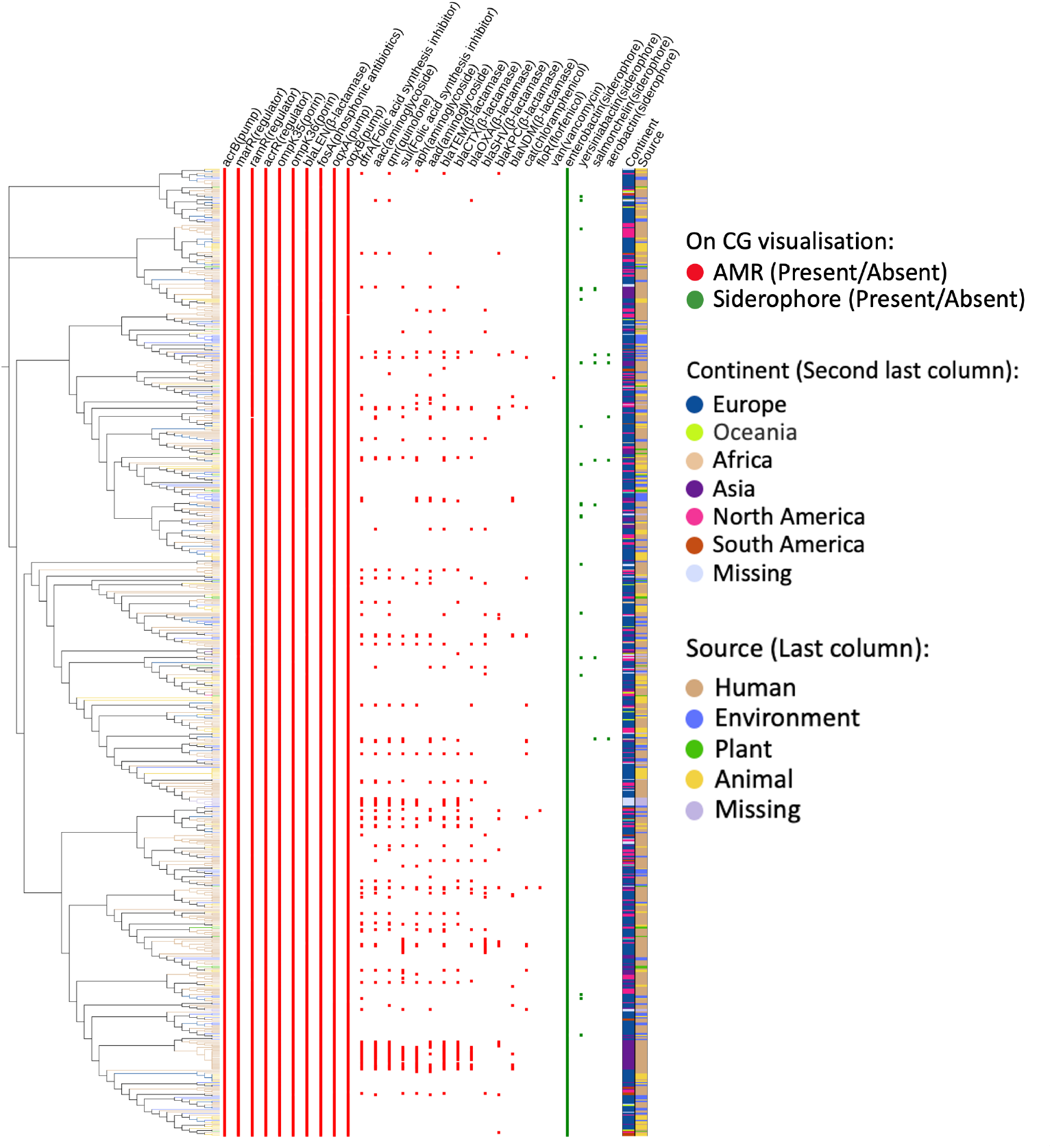
The resistance and siderophore profile of *Klebsiella variicola*. The resistance profile of 715 *Klebsiella variicola* is built using Abricate [37], and the siderophore profile is built by local BLASTP [92]. The resistance and siderophore profiles are visualised on the Core Genome (CG) phylogeny using iTol [67]. The presence of AMR genes is coloured in red, and the presence of siderophores is coloured in green. The geographic locations of the strains are annotated on the second last colour strip, while the host/habitat of the strains are annotated on the last colour strip and the branch colour of the core genome tree.

Interestingly, comparative analysis of AMR gene content across continents reveals that European *K*.*var* strains generally contain fewer AMR genes, while Asian strains exhibit a higher AMR gene count (including *bla*_NDM_ and *cat*) relative to strains from other continents (Supplementary Table 4).

We next aimed to determine whether specific AMR genes were predominantly associated with human isolates. The ortholog groups present in less than 10% of the strains were excluded in GWAS analyses to reduce the multiple test burden and false negatives. Thus, after identifying AMRs with Abricate [37], which provides higher accuracy, we utilised Fisher’s exact test and analysed the AMR genes for whether they are enriched in the human strains (Table 2). As detailed in Table 2, we identified 12 AMR genes significantly associated with the human host which include the *bla*_CTX-M_ (mainly *bla*_CTX-M-15_), *bla*_TEM_ (mainly *bla*_TEM-1_) (encodes class A extended-spectrum *β*-lactamase), *bla*_KPC_ (mainly *bla*_KPC-2_) (encodes carbapenemase), *bla*_OXA_ (mainly *bla*_OXA-1_) (encodes class D *β*-lactamase), *bla*_SHV_ (mainly *bla*_SHV-12_ and *bla*_SHV-1_) (Table 2). With the exception of *bla*_SHV_, which is located on the chromosome, *bla*_OXA_, *bla*_KPC_, *bla*_CTX-M_, and *bla*_TEM_ were located on the plasmids, while *aad* (aminoglycosides) and *cat* (chloramphenicol) are located on both the chromosome and plasmid. In fact, GWAS analyses [32] of human associated ESBL-carrying (n = 56) and non-ESBL carrying *K*.*var* strains (n = 273) confirmed that the ESBL containing strains are more enriched in resistance genes including *bla*_TEM_, *bla*_CTX-M_, *tet*, and *cat*.

**Table 2:** Antibiotic resistance genes associated with the human host in *Klebsiella variicola*. The associations are identified using 441 human strains and 248 non-human strains. The resistance gene name and class are presented in the first and second columns. The *p* values from Fisher’s exact test are shown in the third column, and odds ratio values are presented in the fourth column. The genomic location column refers to the genomic location of this gene, where both indicate presence in both the chromosome and plasmid contigs.

| AMR gene | Class | Fisher test $p$ value | Odds ratio | Genomic location |
| --- | --- | --- | --- | --- |
| *aac* | Aminoglycosides | 1.186e-11 | 39.28 | plasmid |
| *aad* | Aminoglycosides | 2.942e-06 | 7.210 | both |
| *aph* | Aminoglycosides | 2.559e-05 | 5.472 | plasmid |
| *bla<sub>CTX-M</sub>* | β-lactams | 5.086e-07 | 13.17 | plasmid |
| *bla<sub>KPC</sub>* | β-lactams | 0.0002 | infinity | plasmid |
| *bla<sub>OXA</sub>* | β-lactams | 2.838e-08 | infinity | plasmid |
| *bla<sub>SHV</sub>* | β-lactams | 4.247e-05 | 16.60 | chromosome |
| *bla<sub>TEM</sub>* | β-lactams | 1.211e-07 | 8.612 | plasmid |
| *cat* | Chloramphenicol | 0.0237 | 7.437 | both |
| *dfrA* | Folic acid synthesis inhibitor | 4.067e-06 | 4.680 | plasmid |
| *qnr* | Quinolone | 7.541e-11 | 36.33 | plasmid |
| *sul* | Folic acid synthesis inhibitor | 5.495e-08 | 14.89 | plasmid |

As the AMR genes linked to the human *K*.*var* strains are mostly plasmid-located, we used PlasmidFinder (Version 2.0) [38] to generate plasmid profiles of human and non-human associated *K*.*var* strains. Our analyses showed that despite differences in specific gene carriage, the plasmid profiles for human and non-human *K*.*var* are similar (Figure 5A), where the most prevalent plasmid type in *K*.*var* is IncFIB, followed by IncFII and IncFIA. Interestingly, the prevalent plasmid types in *K*.*var* are also similar to our comparator *K*.*pne* subset (Supplementary Table 1). Given the similarities in plasmid profile, we applied Welch’s unequal variances *t* test and found that human *K*.*var* strains (mean = 1.714) contain significantly more plasmids than non-human *K*.*var* strains (mean = 1.009, *p* value *<* 0.001). As shown in Figure 5B, the four most commonly occurring plasmids, IncFIB, IncFII, IncR, and IncN are linked to higher AMR gene carriage but only in *K*.*var* strains from the human host regardless of geographic location.

**Figure 5.**
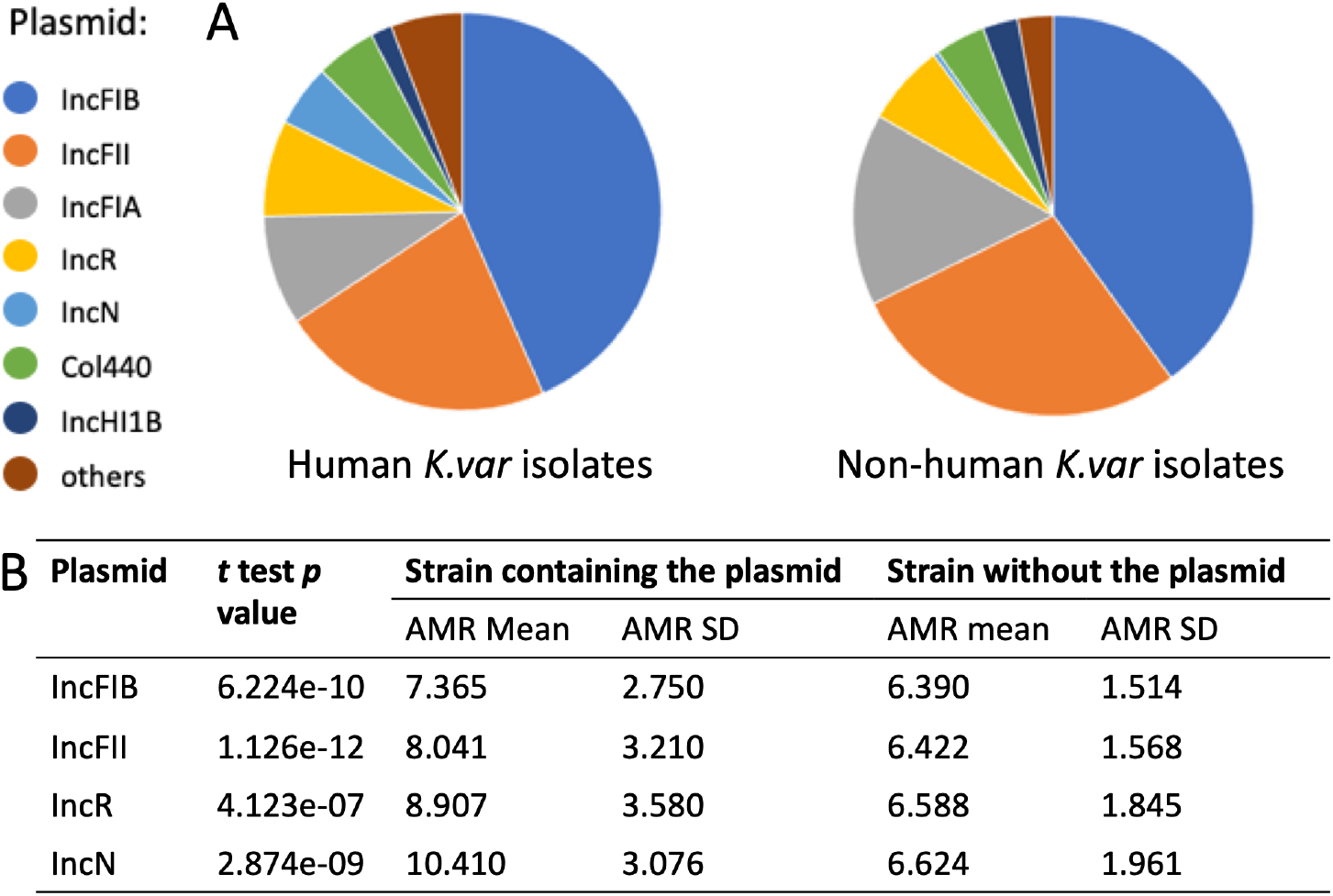
Plasmid Profile of *Klebsiella variicola*. A) **Host/habitat-Based Plasmid Carriage:** The comparison of plasmid carriage between human (n = 441) and non-human (n = 248) strains. The plasmid profile is identified with PlasmidFinder [38]. B) **Plasmids and AMR Gene Carriage:** The plasmids associated with enriched AMR gene carriage in *Klebsiella variicola* (n = 715). In the table, the plasmid type is presented in the first column, and the *t* test *p* values are shown in the second column. The mean and Standard Deviation (SD) values of either strain containing or not containing the plasmid are shown in the following columns correspondingly.

Given that extrinsically acquired AMR genes exhibit clear associations with the human host, we wondered if variations in intrinsic chromosomally located genes which confer antibiotic resistance would follow a similar trajectory. Thus, we constructed gene phylogenies and analysed SNP variations in key intrinsic resistance genes. We based the selection of these intrinsic loci on well-characterised permeability factors including transcriptional proteins (RamR, MarR, AcrR, OqxR, SoxR) where RamR, MarR, AcrR, SoxR, and OqxR function as the cognate repressors of the *ramA, marA, acrAB*, and *oqxAB* genes respectively, also known to regulate efflux (AcrAB, OqxAB) and influx (OmpK35, OmpK36) [39, 40, 41, 42, 43, 44, 45, 46].

The chromosomally located intrinsic OqxR-OqxAB efflux pump locus [47] was present in almost all the *K*.*var* strains, where only two strains from the human host (WUSM KV 35 and 653 KVAR) did not harbour the *oqxA* gene. All SNP variations in the amino acid sequence observed in the dataset for both OqxAB and OqxR (Supplementary Files 1 to 3) were predicted to be neutral by PolyPhen-2 [48].

Our analyses confirmed a trend of increased deleterious sequence variations, as predicted using PolyPhen-2 in *ramR* (*p* value *<* 0.001) amongst human *K*.*var* strains (human = 35, non-human = 2, missing source = 12) (Supplementary Table 5). In contrast, *ompK35* (human = 8, missing source = 5), *ompK36* (human = 16, non-human = 12, missing source = 2), and *acrR* (human = 226, non-human = 171, missing source = 11) harboured deleterious mutations, but these were equivalently found in both human- and non-human strains, suggestive of their broader role in nutrient uptake and cellular homeostasis (Supplementary Table 5). The correlation tests between the intrinsic mutations and plasmid carriage did not produce any significant associations (*p* values: *mrkD* = 0.101, *ramR* = 0.095, *acrR* = -0.303, *ompK35* = 0.192, *ompK36* = -0.071).

We found that 28 of the human *K*.*var* strains in our dataset harbour the *K*.*pne*-associated intrinsic *β*-lactamase *bla*_SHV_ (Figure 4). In previous work [22], *K*.*var* isolates were classed as ‘hybrid’ when at least 3% of the *K*.*var* genome is acquired from another KpSC species isolated from the same hospital and timeframe. Given that the acquisition of *bla*_SHV_ potentially represents a horizontal gene transfer event from *K*.*pne* into *K*.*var*, we wondered whether the *K*.*var* strains which acquired the *bla*_SHV_ gene allele would also qualify as *K*.*var*-*K*.*pne* hybrid strains.

To establish if the *bla*_SHV_-containing *K*.*var* strains would class as hybrid strains, we calculated the ANI of the *K*.*var* strains against both a *K*.*var* (strain: LEMB11, genome assembly: GCF 009648975.1) and a *K*.*pne* reference genome (strain: HS11286, genome assembly: GCF 000240185.1). We selected these reference genomes due to the high quality of the genome assembly which would limit errors in our analyses. Our ANI results show that there were no significant differences in ANI (*p* value = 0.5868) between either the *bla*_SHV_-positive or negative *K*.*var* strains against both the *K*.*var* and *K*.*pne* reference genome. Whilst this result indicates that there is no evidence of large-scale horizontal gene transfer, the fact remains that *bla*_SHV_ is a *K*.*pne*-associated gene, rarely found in *K*.*var*. In fact, the *bla*_SHV_ gene in all 28 of the *K*.*var* strains showed more than 95% identity to the *bla*_SHV_ in *K*.*pne*. Thus, the gene transfer was limited to the *bla*_SHV_-locus. In support of a localized and targeted gene transfer event, we studied the *bla*_SHV_ gene and the capsule locus in a *bla*_SHV_ containing *K*.*var* strain. Using the *bla*_SHV-1_ carrying *K*.*var* strain SPARK 2054 C2, which was co-isolated with a *bla*_SHV-1_ carrying *K*.*pne* strain (SPARK 2054 C1), we show that the ANI of SPARK 2054 C2 (99.11%) and the *K*.*pne* strain SPARK 2054 C1 (94.63%) are not significantly different from the ANI reported for other *K*.*var* and *K*.*pne* strains in our dataset (average ANI with *K*.*var*: 99.03%, average ANI with *K*.*pne*: 94.62%) overall. The capsule types of these strains are reported as KL11 (SPARK 2054 C1, *K*.*pne*) and KL123 (SPARK 2054 C2, *K*.*var*) respectively. In contrast, the *bla*_SHV-1_ gene shared between these strains has 100% identity. Thus, we infer that although *bla*_SHV_ is obtained from *K*.*pne*, the *K*.*var* strains containing the *bla*_SHV_ gene did not undergo extensive genomic recombination. Interestingly, the *K*.*var* strains carrying the *bla*_SHV_ gene contained more plasmidic resistance genes overall (*dfrA, suI, aac*, and *bla*_TEM_) (mean = 10.82) relative to non-*bla*_SHV_ carrying strains (mean = 6.57). Taken together, our data clearly underscore the higher content of AMR gene carriage in human *K*.*var* isolates.

### 2.6 Capsule Diversity in *Klebsiella variicola*

We next addressed trends in established virulence traits in human *K*.*var* to establish host/habitat-dependent capsule specialisation. Capsular polysaccharides (CPS) are recognised as an important virulence factor in *Klebsiella* pathogenicity where KL1 and KL2 are the most prevalent capsule loci in *K*.*pne* [49, 50]. Capsule synthesis in *K*.*pne* is initiated by glucose phosphotransferase WcaJ, or the galactose phosphotransferase WbaP, which links either glucose or galactose to the lipid carrier molecule. These residues are flipped onto the inner membrane by flippase Wzx and polymerized by O-antigen ligase Wzy. The capsule length is regulated by polysaccharide biosynthesis tyrosine autokinase, Wzc, and lastly the mature polysaccharide chain is exported by polysaccharide export protein Wza [51]. Capsule profiles generated using Kaptive [23] show that 13.8% (n = 99) of *K*.*var* strains possess a novel CPS locus, revealing 28 novel capsule loci unique to *K*.*var* (Supplementary Figure 4).

Our analyses also show that *wcaJ* and *wbaP* are found in just 60.84% (n = 435) and 32.17% (n = 230) of *K*.*var* strains (Figure 6A) respectively. Other genes known to be essential in *K*.*pne* for capsule production, such as *wzx* and *wzy*, are absent in 12.45% and 19.02% of *K*.*var* strains (Figure 6A). With the exception of the flanking genes such as *galF, gnd*, and *ugd*, the most prevalent capsule gene in *K*.*var* is *wza*, thus we infer that the combination of the most prevalent 10 genes critical for capsule production in *K*.*var* are *galF, cpsACP, wzi, wza, wzb, wzc, wzy, wzx, gnd*, and *ugd*. Other capsule synthesis genes such as *manCD* and *rmlABCD* were present at similar prevalence in both *K*.*var* and *K*.*pne*. In both mucoid classical *K*.*pne* and hypervirulent *K*.*pne* (Hv*Kp*), the loss of *wcaJ* is linked to the loss of capsule and reduced virulence, underscoring the role of this initiating phosphotransferase [52, 53] in capsule synthesis. However, *K*.*var* strains such as the Bangladeshi outbreak isolates [10], which carry the ‘KL34 variant’ capsule locus (Figure 6C), lack *wcaJ, wzc*, and *wzy* genes but still produce a significant capsule suggesting the biological function of *wcaJ* in capsule production is different in *K*.*var* relative to *K*.*pne* or is subject to genetic regulation by other factors.

**Figure 6.**
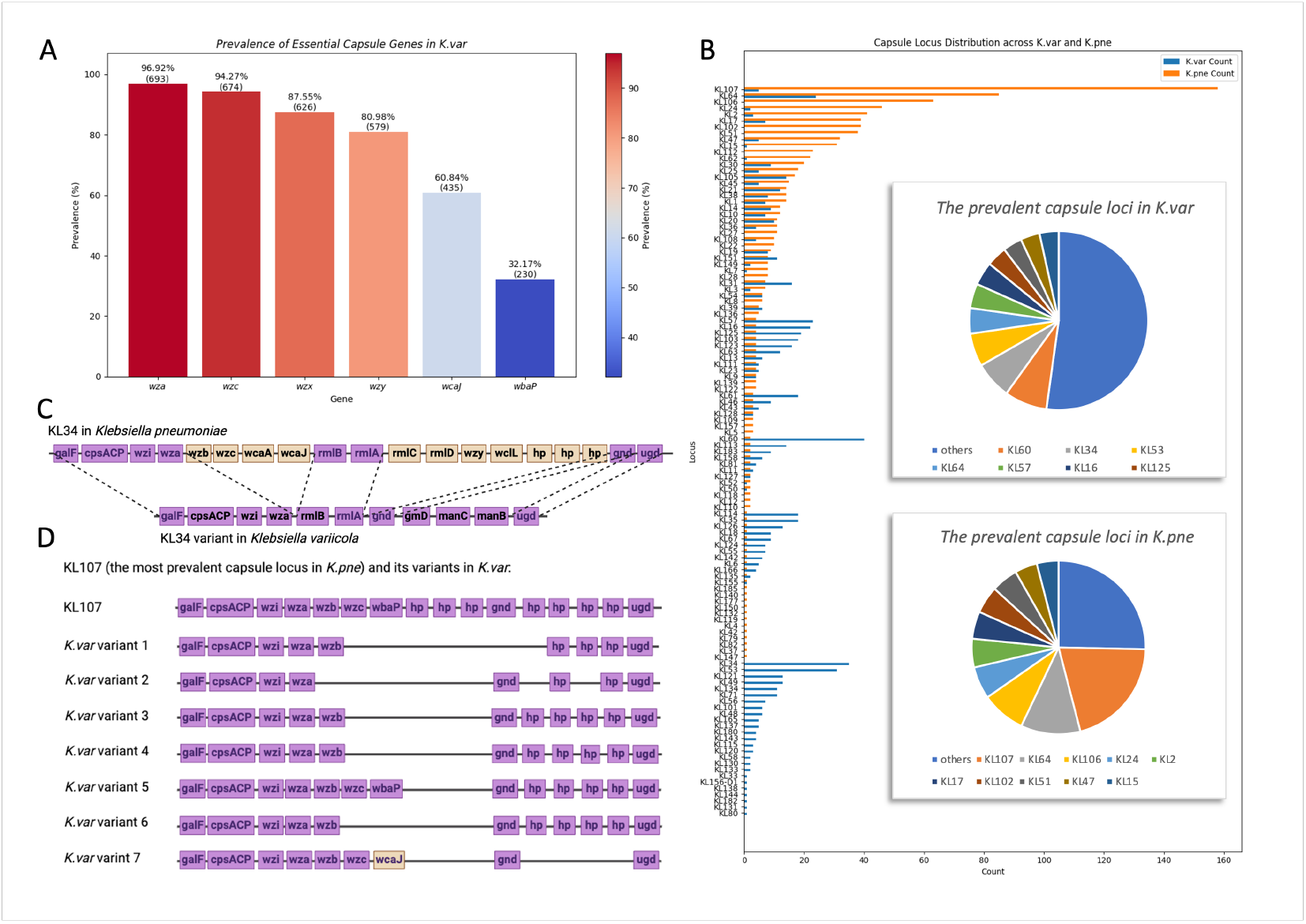
Comparative Analysis of Capsule Loci in *Klebsiella variicola* and *Klebsiella pneumo-niae*. The capsule locus of 715 *Klebsiella variicola* and 1000 *Klebsiella pneumoniae* strains was identified using Kaptive [23]. A) **The heatmap prevalence of inferred essential capsule synthesis genes in *Klebsiella variicola***. B) **The comparison of overall capsule loci in *Klebsiella pneumoniae* and *Klebsiella variicola***. The capsule locus bar plot is ranked in descending order in *Klebsiella pneumoniae*. The most prevalent 10 capsule loci in *Klebsiella variicola* and *Klebsiella pneumoniae* are visualised by pie charts. C) **Novel Capsule Loci in Bangladeshi Strains:** Novel capsule loci specifically found in Bangladeshi neonatal *Klebsiella variicola* strains (KL34 variant) and their comparison with KL34 in *Klebsiella pneumoniae*. Note that ‘hp’ represents hypothetical proteins. D) **A comparison of genes in the most prevalent capsule loci in *Klebsiella pneumoniae* (KL107) and its variants in *Klebsiella variicola***. The matched or differing genes in 6B and 6C are coloured in purple or yellow. Capsule locus images were generated with Biorender.com.

The capsule regulators such as KvrA, KvrB, KbvR, RfaH, and RcsB were highly conserved in *K*.*var* and the sequences showed near uniform alignment with *K*.*pne* (Supplementary Files 4 to 8). The *rmp* genes were only found in 6 *K*.*var* strains (human = 5, missing source = 1) (Supplementary Table 6). A comparative analysis of the ESBL (n = 56) and non-ESBL (n = 273) containing human strains shows that the most prevalent capsule locus carried by non-ESBL containing strains is KL60 (n = 30, percentage = 10.99%). In contrast, the ESBL containing human strains carry diverse capsule types, including the KL34 variant (n = 17, percentage = 23.21%), KL105 and KL123 (n = 4, percentage = 7.14%), KL14 (n = 3, percentage = 5.36%) and KL60, KL31, KL10, KL58, KL126, KL108, KL183, KL56, and KL125 (n = 2, percentage = 3.57%).

Overall, we found that whilst 80 capsule loci are shared between *K*.*var* and *K*.*pne*, 25 loci were unique to *K*.*var* and 12 loci are predominantly found in *K*.*pne* (Figure 6B). Interestingly, the most prevalent capsule loci in *K*.*pne*, such as KL1 and KL2 [49, 50], are rare in *K*.*var* (Figure 6C). Our analyses identified 64 capsule loci present in less than 1% of our *K*.*var* database, implying that *K*.*var* may have acquired these loci from *K*.*pne*. For example, KL107, which is largely present in *K*.*pne* (158/1000), is found in only 5 *K*.*var* strains with coverage identity of 99.99%, suggesting capsule locus acquisition from *K*.*pne* to *K*.*var*. Conversely, KL60, which is more prevalent in *K*.*var* (40/715) relative to *K*.*pne* (2/1000), is found with 100% coverage identity in *K*.*pne*, indicating potential acquisition from *K*.*var* by *K*.*pne*. These observations suggest the bidirectional acquisition of capsule loci between *K*.*var* and *K*.*pne*.

### 2.7 Other virulence genes

Finally, our assessments of the *mrkD* gene, known for its role in type III fimbriae and biofilm formation, showed several deleterious mutations such as T88M and S29T (Supplementary Table 5) which were significantly associated with human *K*.*var* strains (*p* value = 0.0119). Similarly, higher levels of siderophore gene carriage are found in strains isolated from human infections, with enterobactin being the most highly conserved followed by yersiniabactin (Figure 4). However, we observe limited evidence of convergence of resistance and siderophore gene carriage in *K*.*var* as strains which carried more AMR genes did not always have a higher siderophore gene content. The AMR count of siderophore-positive strains (mean = 6.696) is similar to the siderophore-negative strains (mean = 6.566).

## 3 Discussion

Using the largest *K*.*var* database analysed to date, we show that *Klebsiella variicola* is broadly and widely distributed across multiple sources (Figure 1A). The star-like population structure attests to this distribution and confirms diversification from a single non-human common ancestor over a 1000-year time scale (Figure 2A and 2B). This multisource dissemination contrasts findings from a previous study using *K*.*pne* [8] collected within a defined geographic area which shows limited transmission from the environment and animal reservoirs to the human host.

Our analyses demonstrate that this adaptability is underpinned by discernible genetic features which delineate human versus non-human *K*.*var* strains. As a testament to its environmental origins shown by ancestral state mapping, GWAS analyses (Table 1) suggest that non-human associated *K*.*var* have genes which either facilitate environmental survival where the gene attrition noted for these genes specifically *lpxP, chbP*, and *cspB* in human *K*.*var* is a hallmark of adaptation to the human host. In contrast, human *K*.*var* strains are enriched in AMR gene content, AMR gene diversity (Figure 3) and mutation in intrinsic AMR genes e.g., *ramR* (Supplementary Table 5), which results in phenotypes associated with reduced antibiotic susceptibility and increased immune evasion [42, 54, 55]. The higher AMR gene content and diversity are inferred to be largely driven by a combination of plasmid gene acquisition and mutations in intrinsic chromosomal loci where correlation tests suggest that acquisition and mutation events likely occur independently which highlight multiple routes to the emergence of antibiotic resistance in *K*.*var*.

The clear differences in the levels of antibiotic resistance gene carriage in human associated *K*.*var* underscore the potential for gene exchange between *K*.*var* and *K*.*pne*. Firstly, similar plasmid types are found in both species clearly facilitating the cross-transfer of these elements between these species [4]. Secondly, localised HGT events such as the acquisition of the *K*.*pne*-associated chromosomal *β*-lactamase *bla*_SHV_ were observed. Interestingly, we did not identify the presence of the LEN *β*-lactamase in our comparator *K*.*pne* dataset or in online analyses of publicly available *K*.*pne* sequences [56, 57, 58] suggesting that *K*.*var* is a better recipient as opposed to a donor of genetic material which is consistent with the higher rates of recombination between *K*.*pne* and *K*.*var* (8.69%) rather than within *K*.*var* itself (1.57%).

In *K*.*pne*, there is evidence that resistance gene acquisition (e.g. *bla*_KPC_, *bla*_TEM_, *bla*_OXA_, and *bla*_NDM_) is linked to antibiotic use in human *K*.*pne* infections [59] and progression from colonisation to infection [60]. Our analyses (Table 2) show that *K*.*var* follows the same trajectory.

We (Figure 4, Table 2) and others [22, 8] show that *K*.*var* co-isolated with *K*.*pne* can acquire both resistance and capsular genes. Accordingly, capsular profile analyses of *K*.*var* strains which show differential carriage rates suggest that the genetic acquisition of certain capsular types e.g., KL107 and KL60 can occur between *K*.*var* and *K*.*pne* in a bi-directional manner. The capsular polysaccharide locus in *K*.*var* is notably diverse relative to *K*.*pne* (Figure 6B). Whilst the knowledge of the exact impact of these variants on capsulation remains limited, it is important to note that the key capsule synthesis genes such as *wcaJ, wzc*, and *wzy*, shown to be critical for capsule production in *K*.*pne* [51, 52, 53], are not critical for capsulation in *K*.*var* [10]. This contrasts with the high levels of conservation seen for capsule transport systems (*wza*) and highlights that the regulation of capsule synthesis and production is likely different in *K*.*var*. Importantly, the most commonly targeted capsular types in vaccines (KL1 and KL2) are nearly absent in *K*.*var*, limiting the use of *Klebsiella* vaccines against *K*.*var* [61, 62].

We note that there is limited evidence of convergence of both antibiotic resistance and virulence genes in human *K*.*var* isolates. For example, the acquisition of resistance genes in *K*.*var* is not matched by capsular specialisation as seen in certain epidemic lineages in *K*.*pne* e.g., ST11 [63]. In fact, comparative analysis of ESBL and non-ESBL producing human strains do not demonstrate capsule specialisation as ESBL producers have diverse capsule types. Nonetheless, human *K*.*var* isolates showed increased deleterious mutations in virulence loci (Supplementary Table 5) e.g., *mrkD* and siderophore carriage (Figure 4) which promote biofilm formation and increased virulence in vivo [54, 64, 55].

Taken together, our work provides a mechanistic basis for the emergence of *K*.*var* as an emerging human pathogen under antibiotic pressure with genetic and spacial proximity to *K*.*pne*. This, in combination with the lack of ecological niche separation of *K*.*var* isolates, highlights the potential for the spillover of human *K*.*var* into the broader environment and for *K*.*var* isolates within the broader environment to adapt to human infection.

Our work cautions against the application of a one-size-fits-all approach in the genomic analyses of KpSC and underscores the distinctive genetic attributes and transmission routes of *K*.*var*, which stress the emergence of this species as a human pathogen to note with direct implications in the One Health framework.

## 4 Materials and Methods

### 4.1 Data availability

In this work, we acquired 723 *Klebsiella variicola* genomes, which contain 715 *Klebsiella variicola* ssp. *variicola* (*K*.*var*) and 8 *Klebsiella variicola* ssp. *tropicalensis*. This collection of *K*.*var* includes 417 strains accessible via the National Center for Biotechnology Information (NCBI) in September 2022, 279 strains provided by Edward Feil [8], 3 unpublished genomes from Refath Farzana’s study in 2019, including those published on NCBI [10], 3 strains from Sands et al.’s study [65], and 13 *K*.*var* strains sequenced using Illumina short-read sequencing provided by Paul Higgins [66]. We analysed a total of 715 *Klebsiella variicola* ssp. *variicola* genome sequences for core genome phylogeny and resistance and virulence profiles, while strains missing host/habitat information (n = 26) were excluded from comparative analyses. The corresponding metadata for these isolates, including host/habitat, geographic location, and collection date, were sourced from the NCBI BioSample database. We merged the animal, plant, and environmental strains into non-human host/habitats (n = 248) to complement the human host (n = 441). Geographic locations were classified into continents. For transcontinental countries, e.g., Turkey and Russia, the continent of the strain is classified by the exact collection location in the country. The collection dates were categorised by year. Additionally, we built a representative dataset of *Klebsiella pneumoniae* (*K*.*pne*) for comparative analysis by randomly selecting 1,000 genome assemblies from the publicly available *K*.*pne* assemblies on the NCBI database. The sampling of the *K*.*pne* database was regardless of the ST and any metadata on the BioSample page. The details of the datasets for *K*.*var* and *K*.*pne* are presented in Supplementary Table 1 and also made available in Edinburgh DataSync (https://datasync.ed.ac.uk/index.php/s/mZRtOKr6hfq7FSD and https://datasync.ed.ac.uk/index.php/s/tr9L9csL3S31qjk).

During the writing of this manuscript, we discovered further *K*.*var* genomes in the NCBI database. We included a further 167 strains from NCBI, resulting in an expanded database of 882 strains (Supplementary Figure 2). These additional genomes have now been included to generate an updated core genome phylogeny tree (Supplementary Figure 5). To demonstrate the robustness of using the smaller *K*.*var* dataset (n=715) in our downstream analyses, we note that the increased number of strains does not alter the pan-genome structure, and importantly, convergence was observed in the number of unique genes (Figure 1B), indicating that this subset (n=715) can sufficiently represent the genetic diversity of *K*.*var*.

### 4.2 Population structure and pan-genome analysis

For the single nucleotide polymorphism (SNP)-based core genome phylogeny of *K*.*var*, we applied ParSNP [25] and iTOL for phylogeny construction and visualization [67]. The recombination-free core genome alignment from ParSNP was converted using Harvesttools [25], and Seqret [68] was used to generate the Nexus (.nex) alignment file. Genome annotation was performed using Prokka with the genus setting as ‘*Klebsiella*’ [69]. For building the pan-genome of *K*.*var*, we employed Roary [70], opting to prevent paralog splitting (-s) and increasing the cluster number setting (-g) to 60,000. This adjustment was necessary to fit the size of our dataset, which exceeded the capacity of Roary’s default cluster number (50,000). The genes of specific interest, including the *nif* and the capsule regulator genes, e.g., *kvrA, kvrB*, were extracted using Roary [70]. Then, multiple sequence alignment was performed using MAFFT [71]. The gene phylogenies were constructed using IQ-Tree [72], and the mutations were predicted using PolyPhen-2 [48]. *K*.*var* Multi Locus Sequence Typing (MLST) profile was generated using Kleborate [23] and MLSTKv(https://mlstkv.insp.mx/) [21], which is a *K*.*var* specific MLST. PHYLOViZ 2.0 [73] was then applied for globally optimized eBURST analysis and visualization. To assess the congruence between the core genome phylogeny and the MLST, we calculated Baker’s Gamma Index [24] as implemented in the dendextend [74] R package.

### 4.3 Statistical analysis of metadata independence

To test if the metadata of *K*.*var* were independent of each other, we conducted a Generalized Linear Model (GLM) analysis in R [75], specifically focusing on host/habitat and continent annotations. We clustered the strains into ‘human’ or ‘non-human’ sources, and the continent category into ‘European’ and ‘non-European’. We applied the GLM to both the simplified and the original metadata sets.

### 4.4 Defining a type *K*.*var*

We used the gene presence and absence matrix generated by Roary [70] to compute Euclidean pairwise distances, utilising the SciPy library [76] within Python. These pairwise distances for the *K*.*var* dataset served as the basis for Multi-Dimensional Scaling (MDS) analyses, as implemented in the stats package in R [75]. Within this MDS framework, the ‘type’ *K*.*var* isolate is defined as the one nearest to the centroid of all *K*.*var* isolates [77].

### 4.5 Average nucleotide identity

The Average Nucleotide Identity (ANI) was determined using FastANI [78] to analyse the genomic similarities within *Klebsiella* strains. We utilised the NCBI reference genomes for *Klebsiella pneumoniae* (*K*.*pne*, GCF 000240185.1) and *Klebsiella variicola* (*K*.*var*, GCF 009648975.1) to ensure the reliability of our ANI calculations. To reduce imbalance and random variation in the dataset, we employed over- and under-sampling of 100 samples for strains with and without the *bla*_SHV_ gene ten times in our comparison. The *t* test was then applied to assess the statistical significance of the ANI differences observed in these re-sampled groups. The *p* value threshold is 0.05.

### 4.6 Recombination

For recombination analyses, we first constructed a core-genome alignment with ParSNP [25] again, specifically opting not to exclude recombination events. This alignment was then converted into FASTA format using Harvesttools [25]. Subsequently, we employed Gubbins [79] to analyse the alignment, utilising the NCBI reference genomes for *Klebsiella pneumoniae* (*K*.*pne*, GCF 000240185.1) and *Klebsiella variicola* (*K*.*var*, GCF 009648975.1) to identify recombinant sequences. We quantified the recombination rate of *K*.*var* as previously described [80] for *K*.*pne* using the formula:

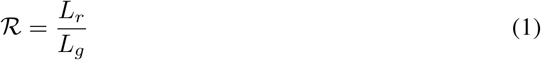

where *ℛ* denotes recombination rate; *L*_*r*_ denotes the total length of all recombination events predicted; *L*_*g*_ denotes genome length.

### 4.7 Ancestral state reconstruction

For ancestral state tracing, we included only those *K*.*var* strains with host/habitat annotation from Supplementary Table 1 (human = 441, non-human = 248). To reduce the overrepresentation of strains from human sources, we grouped the strains from animal, plant, and environmental origins under the ‘non-human’ category. We utilised BayesTraits V3.0.5 [27] to trace the the ancestral state of host/habitat, employing the Maximum Likelihood model in two separate models. The first model did not incorporate any fitted parameters, e.g., *κ, λ*, or *δ*. The second model incorporated the parameter *κ* to assess whether trait evolution is associated with the phylogeny. The log-likelihood for the first model is -839.314049, while for the second model, it is -415.12330. We performed a chi-square test on the Likelihood Ratio (LR) to determine if including *κ* significantly improved the model fit, using a degree of freedom (df) of 1 for a one-sided test according to the manual of BayesTraits [27]:

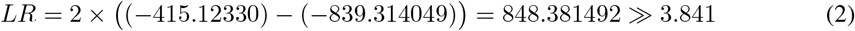

This LR value is significantly higher than the critical value of 3.841, indicating the likelihood of the second model is significantly greater. Therefore, the *κ* parameter offers a more accurate representation of our data, and thus the estimation of *κ* is robust. Accordingly, we chose the *κ*-inclusive model for subsequent analyses.

### 4.8 Time-scaled phylogenetic analysis

For time-measured phylogenetic analysis, a data subset was created with balanced representation of human and non-human isolates for each year in the metadata (Supplementary Table 3). We employed Beast (V2.6.7) [26] for time-measured phylogenetic analysis on this subset. As the sampling of *K*.*var* collection dates clusters largely within the recent five years (2016-2020), the Piqmee package [81] was chosen. We used the default Hasegawa–Kishino–Yano model [82] for nucleotide evolution and implemented a strict molecular clock model sampling 1/5000. Tracer [83] was applied for the examination of the log file parameters. The model parameters converged after 1,000,000 iterations. Nevertheless, we continued the analysis up to 93,665,000 states to confirm the model’s optimal fit. To avoid the risk of getting local optima, we repeated the analyses three times. The repeated runs reached convergence after 20,000,000 states and generated the same results. Next, we consolidated the log and tree files using Log Combiner with default parameters [84]. The maximum clade credibility tree was constructed using TreeAnnotator [84], and finally, FigTree [85] was employed for the visualization and annotation of the time-scaled phylogenetic tree.

### 4.9 GWAS

We conducted Genome-Wide Association Studies (GWAS) to differentiate between human (n = 441) and non-human (n = 248) *K*.*var* strains at the pan-genome level. Our approach included two GWAS methodologies: phylogenetic-based GWAS and pan-genome GWAS. In the phylogenetic-based GWAS, we applied treeWAS [31] in R to assess the statistical association between the phenotype (human/non-human) and genotype. In both approaches, the genes present in less than 10% of the genomes were excluded to increase the statistical power for detecting associations and reducing false negatives. The core genome phylogeny constructed by ParSNP [25] and the pan-genome data generated by Roary [70] were used as the input for identifying significant associations. We further employed both Correlation and Fisher’s exact test, adjusting the *p* values with False Discovery Rate (FDR) [86] and Bonferroni [87] corrections. 0.05 was used as the adjusted *p* value threshold.

For the pan-genome GWAS, we applied Scoary [32] to the Roary-generated data to calculate associations between all genes in the core and accessory genomes and the host/habitat classification (human/non-human).

### 4.10 AMR CARRIAGE

To build the antimicrobial resistance (AMR) carriage profiles for the *K*.*var* database (n = 715), we used ABRicate [37] and then merged the columns representing allelic variations in AMR genes for visualisation. These profiles were then converted into a binary format indicating gene presence or absence and integrated with the core genome phylogeny using iTol [88] for visualisation. Comparison of the human and non-human sourced strains was only done with strains that had a host/habitat annotation (human = 441, non-human = 248). The diversity within the AMR profiles was quantified and graphically represented through the iNEXT package [89] in RStudio. To check the impact of variations in AMR sequences, we utilised PolyPhen-2 [48] to determine whether these genetic alterations in the AMR genes are likely to be neutral or deleterious. We utilised PlasmidFinder (V2.0) [38] to analyse the plasmid carriage profiles within each *K*.*var* isolate. To investigate the association between all plasmids and their corresponding antimicrobial resistance (AMR) carriage, we applied the Welch unequal variance *t* test.

For comparative analyses of resistant and sensitive strains, we excluded the highly multidrug-resistant isolates linked to the neonatal sepsis outbreak in Bangladesh [10], as these strains showed a notably higher number of AMR genes (mean = 16.36) relative to other *K*.*var* strains (mean = 6.73). Extended-spectrum *β*-lactamases (ESBLs) producing human strains are grouped as those which harbour *bla*_CTX-M_, *bla*_OXA_, and *bla*_KPC_ genes [90].

### 4.11 Statistical tests of intrinsic and extrinsic AMR genes

We recorded all variations in amino acid sequences associated with intrinsic and chromosomally located antimicrobial resistance (AMR) genes. The occurrence of these variations was then statistically compared against human/non-human categories using Fisher’s exact test to determine any significant associations. Similarly, for extrinsic AMRs, Fisher’s exact tests were employed to evaluate the relationship between the presence or absence of these genes and the ‘human’ or ‘non-human’ classifications.

We developed a method to determine the genomic location of each AMR gene based on the proportion of core and accessory genes present on the same contig. Usually, the core genes (present in ≥ 95% of the strains) are located on the chromosome, and the accessory genes (present in *<* 95% of the strains) are mostly located on the plasmid. If a contig predominantly consists of core genes, we infer that the AMR gene is located on the chromosome. Conversely, if the majority of the genes on the contig are accessory genes, the AMR gene is inferred to be located on the plasmid. However, if an accessory gene (such as *bla*_SHV_) is located on a contig where the majority are core genes, we categorised this gene as chromosomally located.

### 4.12 Virulence Factors

The capsule locus for each strain was characterized using Kaptive (V2.0) [23]. Loci that did not match any known types in the Kaptive database and were identified as ‘None’ in the output were inferred as novel. Additionally, we extracted the genes in the capsule loci (between the flanking genes *galF* and *gnd*) of KL34 variant *K*.*var* strains and annotated them using Pannzer2 (V2.0) [91] to find potential novel genes. We determined the presence of siderophores by using BLASTP searches [92], with a percentage identity threshold of 95%. The siderophore profiles, including gene presence or absence, were mapped onto the core genome phylogeny and visualized using iTol [88].

### 4.13 Code availability

All relevant Python and R codes are publicly available at https://github.com/JadeWang-0526/K_variicola.git.

## 5 Acknowledgments

JW was supported by the University of Edinburgh and Leiden University PhD studentship scheme. The authors thank Ed Feil for the provision of genome sequences (https://www.ebi.ac.uk/ena/browser/view/PRJEB27342) and Davide Sassera for providing strains SPARK 2545 C1, SPARK 1744 C1, SPARK 2054 C2, SPARK 26 C1, and SPARK 39 C1. The authors thank the Schneiders lab group for critical reading and discussions of the manuscript.

